# Tightly-linked antagonistic-effect loci underlie polygenic demographic variation in *C. elegans*

**DOI:** 10.1101/428466

**Authors:** Max R. Bernstein, Stefan Zdraljevic, Erik C. Andersen, Matthew V. Rockman

## Abstract

Recent work has provided strong empirical support for the classic polygenic model for trait variation. Population-based findings suggest that most regions of genome harbor variation affecting most traits. This view is hard to reconcile with the experience of researchers who define gene functions using mutagenesis, comparing mutants one at a time to the wild type. Here, we use the approach of experimental genetics to show that indeed, most genomic regions carry variants with detectable effects on complex traits. We used high-throughput phenotyping to characterize demography as a multivariate trait in growing populations of *Caenorhabditis elegans* sensitized by nickel stress. We show that demography under these conditions is genetically complex in a panel of recombinant inbred lines. We then focused on a 1.4-Mb region of the X chromosome. When we compared two near isogenic lines (NILs) that differ only at this region, they were phenotypically indistinguishable. When we used additional NILs to subdivide the region into fifteen intervals, each encompassing ~0.001 of the genome, we found that eleven of intervals have significant effects. These effects are often similar in magnitude to those of genome-wide significant QTLs mapped in the recombinant inbred lines but are antagonized by the effects of variants in adjacent intervals. Contrary to the expectation of small additive effects, our findings point to large-effect variants whose effects are masked by epistasis or linkage disequilibrium between alleles of opposing effect.

---

Understanding the genetic architecture of complex traits is a primary goal of quantitative genetics. For more than a century, experimental and theoretical studies have examined the extent to which phenotypes are polygenic and the distributions of effect sizes associated with genomic loci underlying observed variation. Evidence from experimental studies describing quantitative trait variation suggests that polygeny is the norm (Boyle et al., 2017; Mackay et al., 2009; Rockman, 2012). For example, recent analyses of human genetic variation have inferred that the majority of 1-Mb windows harbor variation that affects schizophrenia risk (Loh et al., 2015), and most 100-kb windows affect height (Boyle et al., 2017). These estimates require assumptions about the relationship between allele frequency, effect size, and linkage disequilibrium (reviewed by Yang et al., 2017), and direct assessment of individual polygene effects is difficult in the context of small effects, complex genetic backgrounds, and low minor allele frequencies. Here, we use a classical genetics approach to isolate small genomic intervals and directly assess their effects on complex traits. We ask, for samples of 0.001 of a genome, do the genetic differences between a single pair of chromosomes sampled from a population affect complex trait variation?

The ability to identify loci associated with trait variation is dependent on the constitution of the mapping population and the number of measurements. Two of the most commonly used types of mapping populations are recombinant inbred lines (RILs) and near-isogenic lines (NILs). RILs leverage genotypic replication across random backgrounds, whereas NILs control for background by holding it constant (Doroszuk et al., 2009; Eshed and Zamir, 1995; Keurentjes et al., 2007; Koumproglou et al., 2002; Shao et al., 2010). RILs provide an efficient way to survey the whole genome for loci with significant marginal effects across multiple backgrounds, but those multiple backgrounds also contribute phenotypic variation. Thus, when comparing the phenotype distributions for two genotype classes at a given genetic marker, RILs have abundant variation within each class due to segregating genetic effects. With NILs, those background effects are eliminated, providing greater power to detect differences between focal genotypes. Moreover, when alleles have effects only in certain backgrounds, their marginal effects, detected in RILs, may be quite modest and hard to detect; in NILs, where the background is fixed, epistatic effects are converted to all or none, depending on the background.

We characterized polygene effects directly by using high-throughput phenotyping and high levels of replication to measure heritable differences among almost-identical genotypes. *Caenorhabditis elegans* is well suited for this type of study, as it is small, has a very short generation time, and is naturally inbred (Barriere and Felix, 2005; Frezal and Felix, 2015; Gray and Cutter, 2014). These features allow for relatively quick construction of RILs and NILs, and assaying animals in 96-well plates. Although many traits in *C. elegans* have a simple genetic basis (*e.g*., Palopoli et al., 2008; Ghosh et al., 2012; Zdraljevic et al., 2017), complex traits in *C. elegans* often have polygenic architectures similar to those seen in humans (Noble et al., 2017). We used a sensitizing condition, excess of the metal nickel, to expose variation that might not be visible under favorable laboratory conditions, and we measured individual and population growth rates as a multivariate phenotype. We assayed a large set of recombinant inbred advanced intercross lines (RIAILs; Andersen et al., 2015) from a cross of strains N2 and CB4856, and a collection of NILs carrying small regions of CB4856 donor genome on the X chromosome within an otherwise N2 background (Bernstein and Rockman, 2016) (Figure 1).

**Figure 1.**
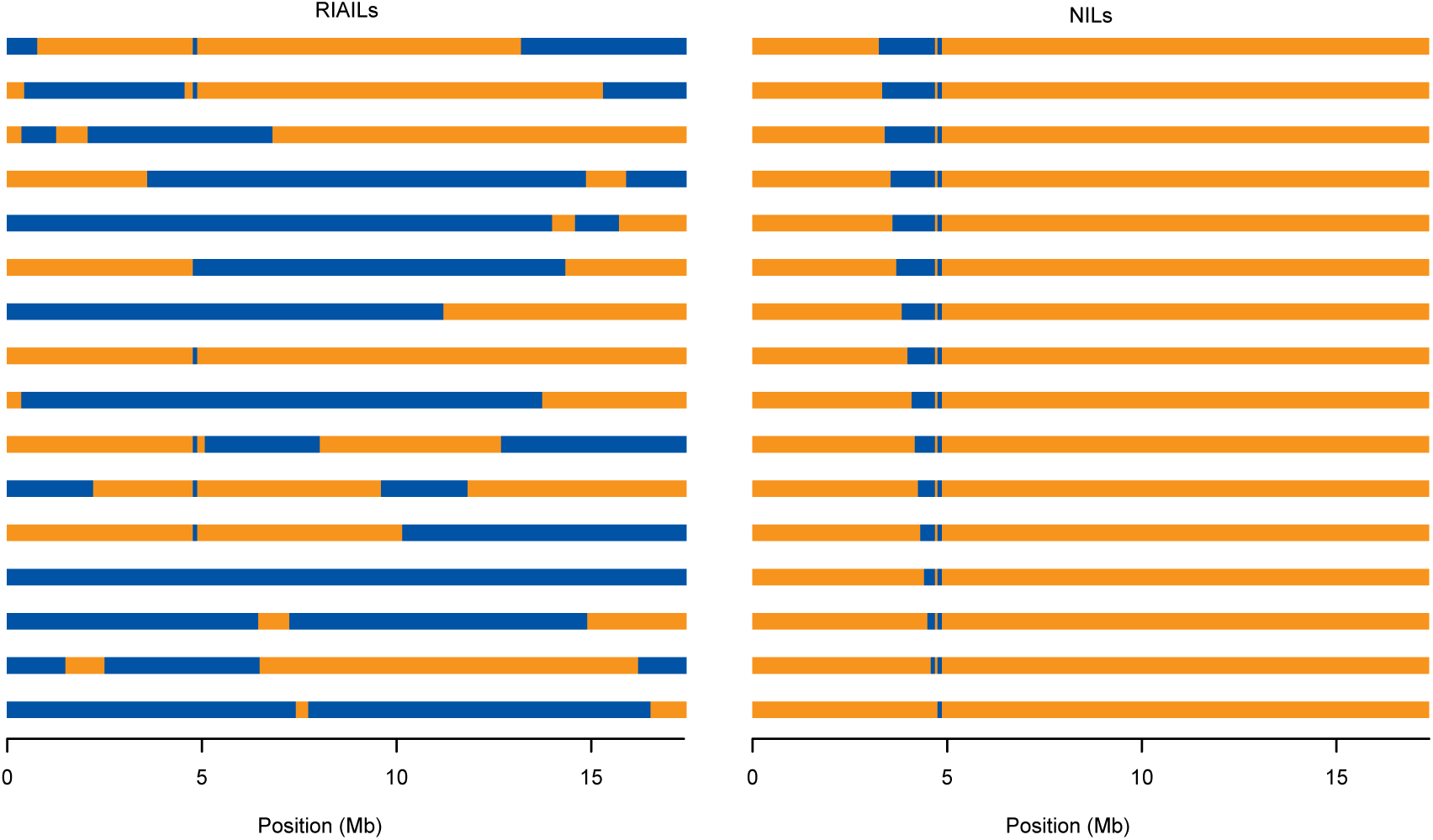
X chromosomes of the two genetic mapping panels. Left: Sixteen of 282 Recombinant Inbred Advanced Intercross Lines, each homozygous for a unique mosaic of N2 (orange) and CB4856 (blue) genome. At any specific marker, approximately half the lines are homozygous N2 and the remainder homozygous CB4856. Right: Near Isogenic Lines derive almost entirely from the N2 background but carry small regions of CB4856 genome with a 1.4 Mb region on the X chromosome. Each CB4856 interval shares a common right end, so that pairs of most-similar strains differ only by 53–148 kb of genome. In both panels, every strain also carries qgIR1, a 110-kb introgression of CB4856 genome at 4.8 Mb; this introgression carries the ancestral allele of npr-1, where N2 carries a laboratory mutation.

## Results

### High-throughput multivariate phenotyping in RIAILs

To characterize the genetic basis for variation in growth and reproduction, we used a previously established pipeline for high-throughput population phenotyping (Andersen et al., 2015; Zdraljevic et al., 2017). We phenotyped N2, CB4856, and 282 RIAILs in liquid cultures containing 350 μM nickel chloride. We chose NiCl_2_ because a preliminary survey of diverse stressors suggested that the left side of the X chromosome carried a QTL for a nickel-by-genotype interaction (data not shown). After placing three L4 hermaphrodites in each well of a microtiter plate and allowing them to mature and produce progeny over four days, we passed each resulting population through a COPAS BIOSORT large-particle sorter (Union Biometrica). The sorter counts the number of animals in each well, and for each animal it measures time of flight, which serves as a measure of body length; extinction, an optical density measure that is a function of body length, size, and composition; and autofluorescence. Body length is a good proxy for the developmental stage of each animal, allowing us to estimate the proportions of different developmental stages, L1, L2+L3, L4, and adult animals, in each well (L2 and L3 larvae cannot be reliably distinguished in this assay). These stage assignments are based on control conditions, and they could be less consistent under nickel stress because growth rate is reduced. The four life-stage proportions within a well must sum to one, so we can use any three proportions to provide a description of the age-structure of the well. We combined offspring number and age-structure into a four-dimensional demographic-state phenotype vector (“demography”) for our subsequent genetic analyses. Assays were completed in 10 blocks (assay days), and trait values were adjusted for assay-day effects by regression (see Methods).

### Demography is heritable

The parental strains, N2 and CB4856, differ significantly in their demographies (MANOVA, *F*_4,15_ = 18.2, p = 1.3 × 10^−5^). N2 has more progeny and a greater proportion of the progeny were measured as adults, consistent with a developmental delay in CB4856 relative to N2.

To determine the genetic contributions to the trait variation we observed, we estimated the broad-sense heritability of demography. We used the phenotypic variance among replicates of each of the parental genotypes to estimate environmental variance, interpreting the excess phenotypic variance among the RIAILs as genetic variance (Brem and Kruglyak, 2005). This method yields estimated broad-sense heritabilities of 0.71 to 0.88 for the four components of the demography phenotype, all significantly greater than zero based on a permutation-derived null distribution (p < 0.01).

### QTL identified among the RIAILs are antagonistic

We performed multivariate QTL mapping to identify regions of the genome that influence demography in the RIAILs. We employed simple multivariate marker regression (Knott and Haley, 2000) on the assay-corrected RIAIL phenotypes, and we used a forward search strategy with a genome-wide p=0.05 permutation-based residual empirical threshold (Doerge and Churchill, 1996). This approach identified seven significant QTL (Figure 2A). The effects of the CB4856 allele at each QTL, projected into bivariate space, are plotted in Figure 2B. Most of the QTL influence multiple aspects of demography, though several are restricted to one or a few trait axes. For example, QTL 5 affects the L2/L3 and adult proportions, but has little effect on L1 proportion or progeny number. QTL 7 influences number of progeny but is nearly orthogonal to the other axes. Moreover, the estimated effects are in both directions for each trait, indicating that parental strains carry mixtures of antagonistic alleles.

**Figure 2.**
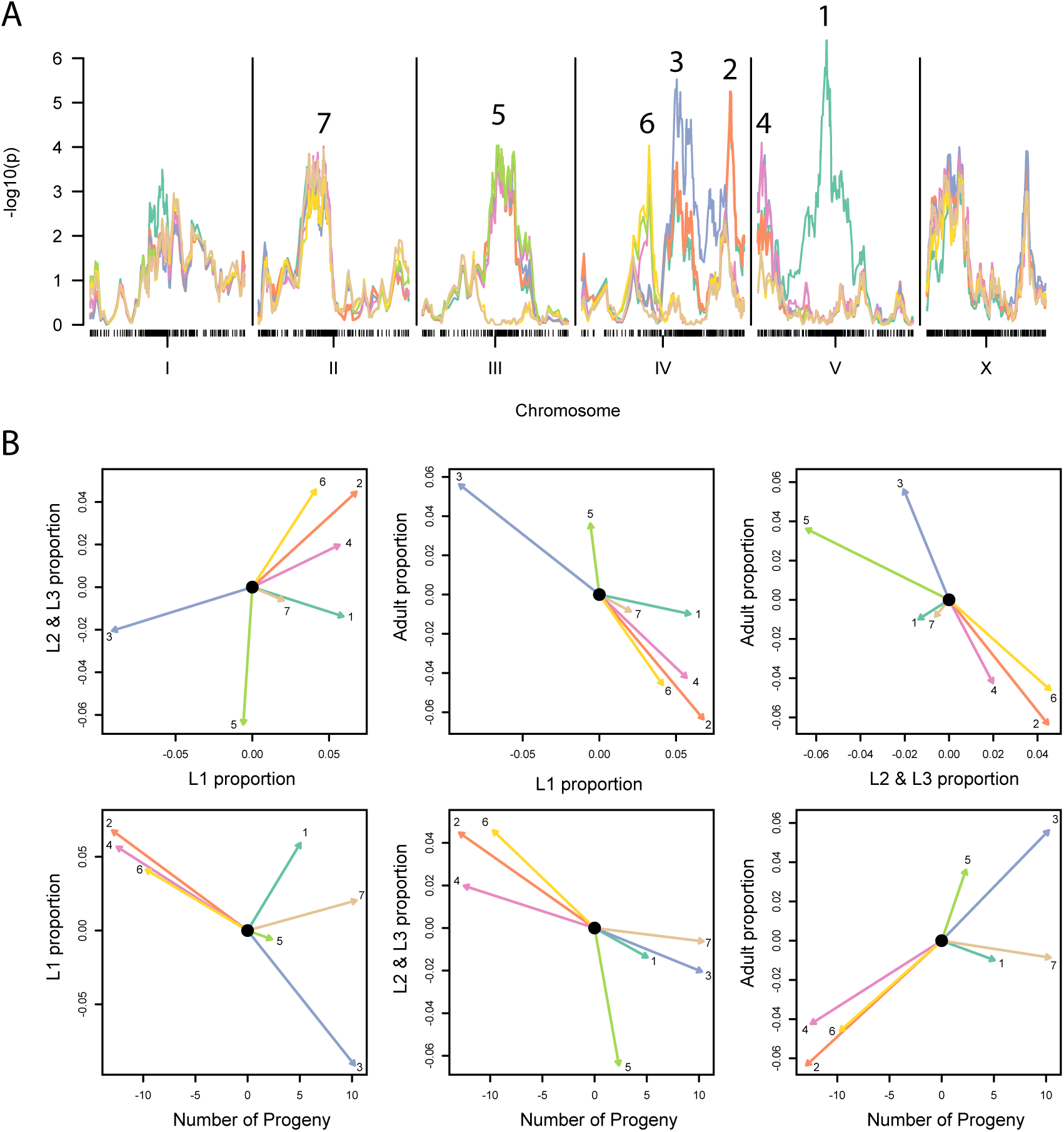
Multivariate demography QTL. A. Multivariate QTL scans with a forward search strategy resulted in a seven-locus genetic model. Test statistic profiles (−log_10_(p)) for seven sequential scans are plotted in different colors, and the QTL retained from each scan indicated by its number. B. Bivariate projections of the QTL effects. Each vector shows the estimated effect of the CB4856 genotype at the indicated QTL, numbered and colored as in panel A.

We found no evidence for pairwise or higher-order epistasis among the detected QTL (p = 0.54 and 0.80 for comparisons to a purely additive model), and the seven-QTL model explains 10, 21, 14, and 31% of the variance in the assay-corrected brood size and L1, L2/L3, and adult proportions, respectively. As the estimated broad-sense heritabilities ranged from 0.71 to 0.88, the great majority of the total genetic variance for each measured trait is not explained by the additive effects of these large-effect QTL.

### Near Isogenic Lines provide a direct test of a polygenic architecture

One possible genetic model for our traits ascribes the unexplained heritable variation to a large number of variants spread across the genome. Under this polygenic model, any region of the genome is likely to harbor phenotypically penetrant variants. We used a panel of 16 Near Isogenic Lines to test 15 consecutive intervals of 53–148 kb (that is, ~0.001 of the 100 Mb genome) spread along a 1.4 Mb region on the X chromosome (Bernstein and Rockman, 2016). Among the RIAILs, this region exhibits elevated but non-significant linkage to demographic traits (Figure 2A). The NILs allow for straightforward tests, comparing two strains that differ only in a single interval, with any phenotypic differences attributable to genetic differences in that interval. These strains simultaneously enable us to control for loci outside the interval (thereby removing a major source of within-marker-class variation), and they expose the variation within the interval that could be masked by tightly linked antagonistic QTL. In total, there are 1,838 SNPs and 635 indels in the NIL interval (Thompson et al., 2015). We assayed growth across three independent experiments. In each of the three experiments, each strain was grown up in five replicate populations over several generations to control for shared environmental effects and then assigned to a random well position that was fixed across 9 to 11 replicate plates within that assay day. In total, we measured NIL demography in 2,312 wells, an average of 145 wells per strain. The design allows us to test the effect of genotype while accounting for variation due to experimental factors. Note that our NIL analysis compares two genotypic classes, each measured approximately 145 times, while in the RIAIL analysis, each marker genotype class is present in approximately 141 (=282/2) strains.

We used univariate mixed-effect models to extract phenotype values (Best Linear Unbiased Predictors) for each of the 12 to 15 replicate populations of each NIL, accounting for variation due to assay day, assay plate within each assay day, and well position. We then applied a fixed-effect multivariate model to estimate each strain’s demography phenotype. As shown in Figure 3, the NILs vary in demography. However, most of the variation is confined to a subspace, as the L1 proportion and L2/L3 proportion are highly negatively correlated. Moreover, the parental NILs, one entirely N2 and the other entirely CB4856 within the NIL region and otherwise genetically identical, have very similar phenotypes and are not significantly different from one another (MANOVA, *F_4,218_* = 2.41, p>0.05).

**Figure 3.**
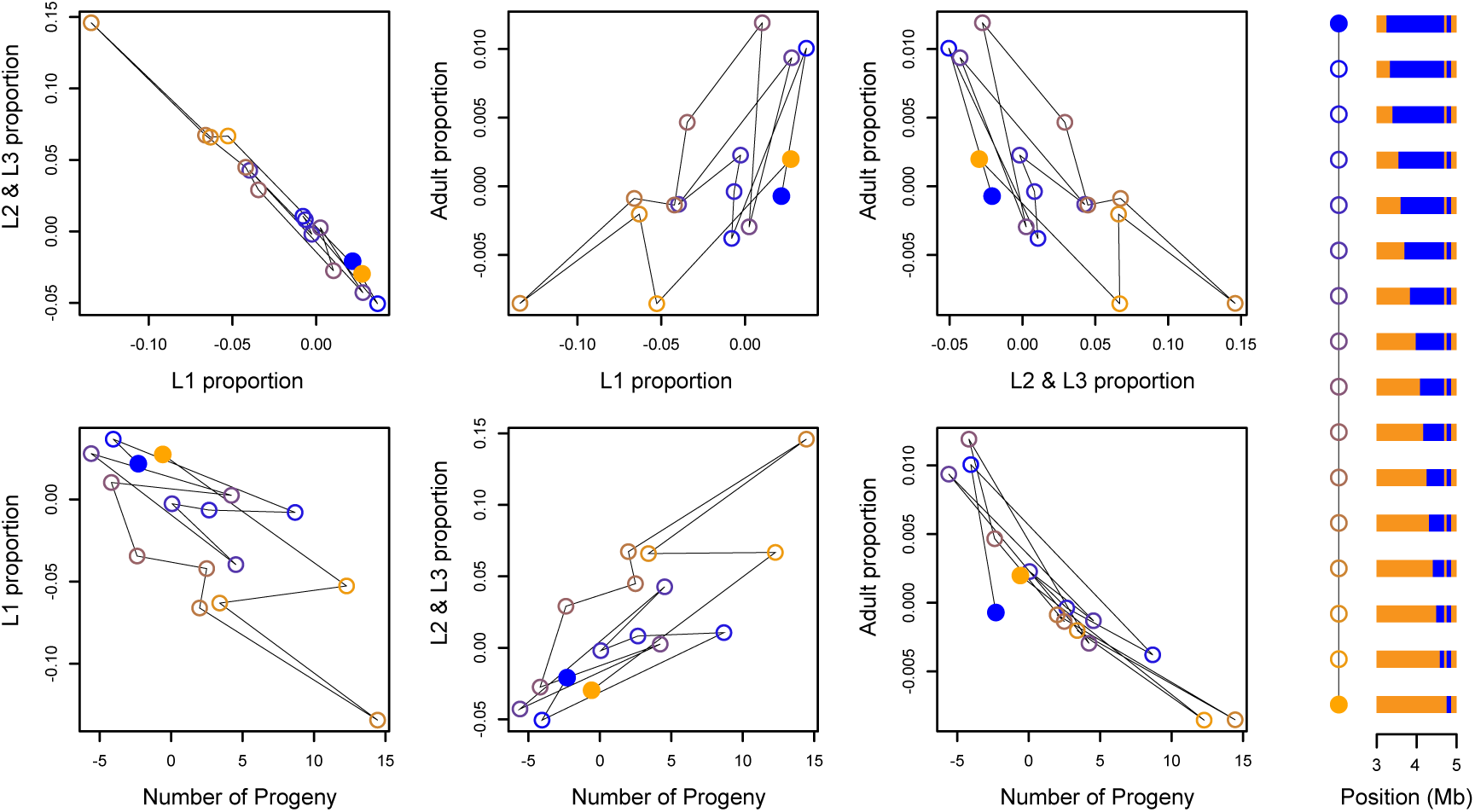
Multivariate phenotypes of 16 Near Isogenic Lines. The filled circles represent the NILs that are entirely CB4856 (blue) or entirely N2 (orange) within the 1.4Mb NIL interval (as shown at right, as in Figure 1). The other NILs are plotted as open circles, connected to one another in the sequence shown at right.

### Multiple QTLs within the NIL interval

We tested whether the demography phenotype of each strain differed significantly from that of the genetically adjacent strain, thereby testing each of fifteen genomic intervals. Eleven of the fifteen intervals contained significant QTLs, at a Bonferroni-adjusted p-value threshold of 0.003 (Figure 4A).

The estimated effects of each NIL interval are plotted in Figure 4B, and they reveal several striking patterns. First, as in the case of the RIAILs, the effects point in both directions for each trait, indicating that the NIL region harbors a mixture of antagonistic QTL. Adjacent intervals often have effects in opposite directions. For example, intervals *l, m, n*, and *o* result in alternate increase, decrease, increase, and decrease in the number of progeny, cancelling one another’s effects. Second, the effects are generally aligned (*i.e*., the vectors are correlated), consistent with the reduced range of variation in certain axes of phenotypic variation. For example, most of the intervals have pleiotropic effects that act antagonistically on L1 proportion and L2/L3 proportion, and all intervals that increase progeny number decrease adult proportion. Nevertheless, as for the RIAILs, some of the NIL QTL affect only one or a few of the phenotypic axes. For example, interval *a* acts almost exclusively on adult proportion, and interval *c* on progeny number. In general, the orientations of the effects are quite different from those observed for the QTL detected in the RIAILs (Figure 2B), indicating a substantially different genetic correlation structure in the two experimental panels. Finally, the magnitudes of the effects in the NILs are large, comparable to those detected in the RIAILs. For example, many of the NIL interval effects change the number of progeny per animal by ten offspring.

**Figure 4.**
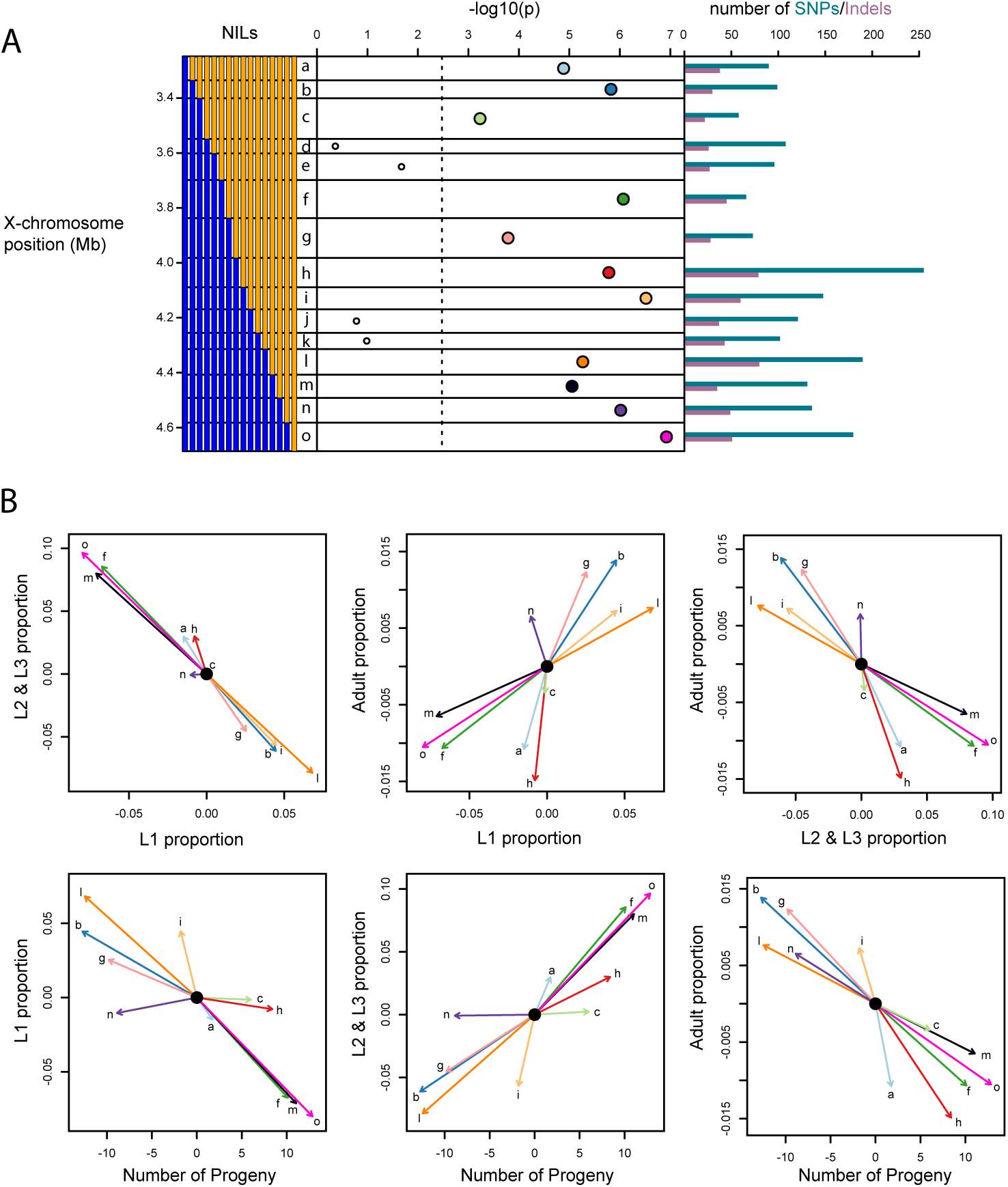
NILs reveal antagonistic QTLs. A. The genotypes of the 16 NILs, at left, define 15 intervals (a-o). By comparing strains that differ only in a single interval, we tested for the effect of the interval on demography, with the results plotted as −log10 of the p-value. The bar plot at right shows the number of SNPs and indels in each interval. Eleven intervals show significant effects. Each significant point is colored to facilitate comparison with panel B. B. Estimated effects of the CB4856 genotype for each significant NIL interval. Note that these effects represent the vectors that connect the NILs in the phenotype space plotted in figure 3.

Simple extrapolation to the 100-Mb genome implies that N2 and CB4856 differ in more than 700 100-kb intervals with significant effects on demography.

The large effect sizes in the NILs raise the question of whether a genome full of such effects is consistent with the variation observed from genome-wide segregation in the RIAILs. We therefore simulated a RIAIL phenotype (offspring number) by assigning effects to a random 700 of the 1454 markers genotyped in the RIAILs, with effects drawn from a normal distribution with the inferred NIL effect-size variance. In 10,000 simulated datasets, the simulated RIAIL variance was on average 10 times greater than the observed RIAIL variance, and never as low as the observed variance. We next simulated effect sizes as uniformly distributed across the range inferred for the NILs, a distribution more similar to the observed pattern. The simulated RIAILs had phenotypic variance on average seven times the observed variance, and again, never as low as the observed variance. These results suggest that either the causal variants do not act additively, or that effects of opposite sign tend to be tightly linked more than expected under a random distribution of effect signs (*i.e*., adjacent antagonistic QTLs cancel one another).

## Discussion

Our NIL-based analysis of small genomic region provides a simple and direct validation of polygeny: most intervals carry segregating variation that affects a complex trait. We used multivariate quantitative genetic analysis to examine the genetic basis for demographic variation in *C. elegans*. Both of our experimental panels, RIAILs and NILs, revealed that two strains, CB4856 and N2, harbor large numbers of allelic differences that affect demographic traits. RIAILs identified significant loci, but most of the heritable variation remained unexplained. When we examined one region of genome more closely, using NILs, we discovered that most small genomic intervals carry phenotypically penetrant variants.

Eleven of fifteen intervals, each ~95 kb, had significant effects on the phenotypes. The focal region of the X chromosome is not particularly noteworthy with regard to trait variation as a whole in these strains (Andersen et al., 2015). Its SNP density is similar to the X chromosome arms as a whole, and the X chromosome arms are considerably less SNP-dense and indel-dense than the autosome arms (Thompson et al., 2015). Our simple extrapolation to more than 700 causal intervals genomewide is likely conservative, given the probability that tightly linked variants within the four non-significant intervals may cancel each others’ effects, causing us to miss them, as we observed for our parent NILs. Moreover, our power to detect very small effects remains quite limited. Projecting to the broader *C. elegans* population, our sample of two strains provides a narrow view of phenotypically relevant genetic variation. N2 and CB4856 differ in our 1.4-Mb focal genomic region by 1,838 SNPs, while a survey of 249 distinct wild isolates (isotypes) identified more than 8,900 single nucleotide variants segregating in the region (Cook et al., 2017) (release 20170531).

Our results contribute to a growing consensus from NIL studies that tightly linked antagonistic QTL (whether additive or epistatic) are a common feature of complex-trait architectures (Gaertner et al., 2012; Glater et al., 2014; Green et al., 2013; Kroymann and Mitchell-Olds, 2005; Shao et al., 2008). Partial selfing may facilitate the evolution of these repulsion-phased complexes, and they may contribute to the widely observed pattern of outbreeding depression in the partially selfing *Caenorhabditis* species (Baird and Stonesifer, 2012; Dolgin et al., 2007; Gimond et al., 2013). Although this pattern may be most common in selfers, it should arise in any system in traits under stabilizing selection, as evidenced by excess repulsion-phase linkage disequilibrium between coding and cis-regulatory variants in humans (Castel et al., 2018). Such linkage disequilibrium provides a simple model for the storage of cryptic genetic variation in natural populations (Kroymann and Mitchell-Olds, 2005; Paaby and Rockman, 2014).

## Materials and Methods

### Strains

We used 282 strains from the Andersen panel of Recombinant Inbred Advanced Intercross Lines (Andersen et al., 2015; Zdraljevic et al., 2017) and 16 strains from the Bernstein panel of Near Isogenic Lines (Bernstein and Rockman, 2016). Construction of the NILs involved use of visible marker mutations, and background mutations from the mutant strains (*fax-1*(*gm83*) and *lon-2*(*e678*); see Bernstein and Rockman 2016) may be present in the NILs. In the construction of both the RIAIL and NIL panels, the N2-like parental strain carried *qgIR1*, an introgression of 110 kb of CB4856 genome on the X chromosome that includes the *npr-1* locus. This gene carries a laboratory-derived mutation in the N2 strain (McGrath et al., 2009), and the *qgIR1* introgression replaces the mutant allele with the ancestral wild-type *npr-1*. Therefore, every NIL and RIAIL carries *qgIR1* and this interval does not contribute to variation within the panels. The RIAIL N2-like parent, QX1430, also carries *ttTi12715*, a transposon insertion in the *peel-1* gene that reduces the paternal-effect incompatibility between it and CB4856 (Andersen). Finally, the N2 strain that was assayed 10 times in parallel with the RIAILs and contributes to our estimate of environmental variance for broad-sense heritability estimation is the reference N2 strain from the *C. elegans* Natural Diversity Resource (Cook et al., 2017).

#### Growth assay conditions

The RIAILs and the NILs were assayed as described previously (Andersen et al., 2015; Zdraljevic et al., 2017). In short, strains were passaged for four consecutive generations to reduce any transgenerational effects from starvation or other stresses. Strains were then bleach-synchronized and aliquoted to 96-well growth plates at approximately one embryo/μl in K medium (Boyd et al., 2012). Embryos were then allowed to hatch overnight to the L1 larval stage. The following day, hatched L1 animals were fed HB101 bacterial lysate (Pennsylvania State University Shared Fermentation Facility) at a final concentration of 5 mg/ml and allowed to grow to L4s over the course of two days at 20°C. Three L4 larvae were then sorted using the Union Biometrica Large-Particle flow cytometer (BIOSORT) into experimental plates containing HB101 lysate at 10 mg/ml, K medium, 31.25 μM kanamycin, and 350 μM nickel chloride dissolved in water. The animals were then grown for four days at 20°C. During this time, the animals matured to adulthood and laid embryos that subsequently hatched and commenced feeding and growing. Prior to measuring the resulting populations’ demographic parameters, animals were treated with sodium azide (50 mM) to straighten their bodies for more accurate length and optical density measurements. Phenotypes that were measured by the BIOSORT include brood size, animal length (time of flight), and animal optical density (extinction). The multivariate phenotype is described in more detail in the results. Data were processed using the R package COPASutils (Shimko and Andersen, 2014), which is available on github.com/Andersenlab/easysorter.

For the RIAIL experiments, each of the 282 RIAILs was assayed once, while N2 and CB4856 were replicated ten times each. These experiments took place over ten assay days. For all analyses of RIAIL data, we used as phenotypes the residuals of a multivariate linear regression of raw phenotypes on assay day, modeled as a factor.

For each of three NIL assay days, each of the sixteen NILs was grown in five independent populations prior to assaying to explicitly control for variation in passaging across strains. The typical NIL was assayed 145–150 times across three experiments. Each strain was assayed in 45–55 wells per assay day. Strain QG2150 was assayed 98 times total due to growth failure on the first assay day. For a given assay day, the position of each strain replicate within the set of nine to eleven plates for that day was randomized.

#### Statistical Analyses

We performed all statistical tests and analyses in R (R Core Development Team, 2017), as described below. Fixed-effect multivariate analyses used the R package *car* (Fox and Weisberg, 2011), and mixed-effect models used the package *lme4* (Bates et al., 2015). The raw data for the RIAILs and NILs are provided as supplements (Supplemental Files S1-S2), and an annotated reproducible pipeline for all analyses is present in Supplemental File S3.

#### Broad-sense heritability in RIAILs

We estimated broad-sense heritability using a multivariate extension of the pooled-variance approach (Brem and Kruglyak, 2005). We compared the phenotypic variance among the RIAILs to within-strain environmental variance, the latter using the pooled covariance estimator for the ten replicates of each parental strain (N2 and CB4856). To assess significance, we performed the same analysis on 100,000 datasets with permuted strain labels, such that ten wells were assigned the N2 label and ten the CB4856 label, with the rest assigned the RIAIL label. All four traits had significant broad-sense heritabilities at p < 0.01.

#### Linkage mapping in RIAILs

We performed multivariate marker regression (Knott and Haley, 2000). The model is **Y** = **X*B*** + **E**, where **Y** is the 282 × 4 matrix of RIAIL phenotypes, **X** is a 282 × 2 design matrix, with 1s in the first column and marker genotypes (0 or 1) in the second, and ***B*** is the 2 × 4 matrix of effects, with intercepts in the first row and marker effects in the second. **E** is the multivariate normal residual error. We fitted this fixed-effect model for each marker using R/qtl (Broman et al., 2003), calculating a p-value for the marker from an approximate F statistic based on the Pillai-Bartlett test statistic. We performed forward selection with a residual empirical threshold to build a QTL model (Doerge and Churchill, 1996). After a marker regression scan, we tested whether any marker exceeded a genome-wide empirical significance threshold of p = 0.05 by analyzing 1000 datasets with permuted strain labels. The top marker genome-wide that passed the empirical threshold was then incorporated into a genetic model of the trait vector, and we repeated the scan and permutations on the multivariate residuals of that model. We continued these cycles of scans and permutations until no additional markers exceeded the relevant residual empirical threshold. The final model includes seven QTL (*i.e*., the design matrix includes eight rows), and we report the variance explained by the model for each trait as the trait-wise multiple *r*-squared from the multivariate model. We tested for epistasis among the seven QTLs using approximate *F*-statistics based on the Pillai-Bartlett trace to compare an additive model and one with all pairwise interactions, and then an additive model and one with all possible interactions.

#### Analysis of Near-Isogenic Lines

We accounted for experimental variation in our measures of NIL demography by treating assay day, assay plate, well position, and biological growth replicate as random effects in univariate analyses of each trait. Each strain was measured in 12 to 15 growth replicates, and we used the Best Linear Unbiased Predictor for each of the 237 growth replicates as the phenotype corresponding to that replicate. To test for the effect of each interval on the multivariate demography phenotype, we compared two fixed-effect multivariate models of the 237 growth-replicate phenotypes. For a given pair of most-related strains, we first modeled the entire data set including strain as a fixed effect with 16 distinct levels, and then we fitted a nested model in which a single pair of most-similar strains are treated as a single genotype. We compared these models using the Pillai-Bartlett test statistic to obtain p-values for the degree of improvement from the merged model to the complete model. We used this same approach to test for a significant difference between the two parental NILs.

## Acknowledgments

This work was supported by NIH R01GM089972 and R01GM121828 (MVR), and funding to ECA from an NIH subcontract (GM107227), the Chicago Biomedical Consortium with support from the Searle Funds at the Chicago Community Trust, and an American Cancer Society Research Scholar Grant (127313-RSG-15-135-01-DD). Support for SZ came from the NIH Cell and Molecular Basis of Disease training grant (T32GM008061).

## Supplementary Files

File S1. RIAIL data

File S2. NIL data

File S3. R code for reproducible analyses.

